# Postnatal Development Shapes the Cardiac Response to Milrinone

**DOI:** 10.64898/2026.09.17.750727

**Authors:** Shatha Salameh, Emily Boozell, May Rajtboriraks, Fiona Mesfin, Anika Haski, Luther Swift, Nikki Gillum Posnack

## Abstract

**Background:** Milrinone, a phosphodiesterase-3 (PDE-3) inhibitor, is widely used to improve cardiac output in pediatric and adult patients. Yet, developmental differences in myocardial responsiveness to milrinone remain incompletely understood. In this study, we examined the impact of postnatal maturation on the acute cardiac effects of milrinone using an intact guinea pig heart model.

**Methods:** Neonatal (0-2 days), juvenile (4-10 days), and adult (> 6 months) guinea pig hearts were excised, Langendorff-perfused, and cardiac metrics were evaluated under basal conditions and in response to acute milrinone treatment (15 minutes, 10 and 100 μM sequential concentrations). Pseudo-electrocardiograms were recorded continuously and left ventricular pressure measurements were performed under sinus rhythm and in response to external pacing. Post-rest potentiation was used to assess contractile reserve, and optical action potentials and calcium transients were recorded.

**Results:** Baseline left ventricular developed pressure (LVDP), action potential duration (APD), and calcium transient duration (CaD) increased with postnatal maturation, consistent with developmental cardiomyocyte remodeling and refinement of excitation-contraction coupling. Milrinone increased heart rate in all age groups, with the greatest chronotropic response at 100 μM. Milrinone also increased ventricular contractility and relaxation across developmental stages, but the magnitude of the response varied with age and was attenuated during high frequency pacing. Adult hearts had the greatest increase in LVDP during sinus rhythm and robust post-rest potentiation, which were less pronounced in neonatal hearts. APD and CaD were shortened in neonatal and adult hearts, but were minimally affected in juveniles, indicating a non-linear developmental pattern.

**Conclusions:** Milrinone exerts positive chronotropic, inotropic, and lusitropic effects throughout development, but the magnitude and frequency dependence of these responses vary with age. These findings suggest that postnatal maturation influences the cardiac response to PDE3 inhibition. These developmental differences highlight the importance of considering age as a biological factor in pediatric drug selection and dosing.

## INTRODUCTION

Congenital heart disease is the most prevalent birth defect affecting approximately 1 out 100 of births worldwide^1^. More than 20% of congenital heart disease patients require corrective heart surgery early in life^2,3^, which is associated with postoperative complications including cardiac arrhythmias and low cardiac output syndrome^4–8^. Antiarrhythmics, inotropes, and vasopressors are frequently employed to stabilize pediatric patients during their postoperative recovery^5,9–11^ – yet only 4% of clinical trials have studied cardiac-specific medications in younger populations^12^. Accordingly, inotropic agents are frequently administered off-label to neonatal, infant, and pediatric cardiac patients, with drug selection and dosing regimens largely extrapolated from adult clinical data. Milrinone, a phosphodiesterase (PDE) 3 inhibitor, is one of the most commonly prescribed inotropic agents (up to 97.9% of patients)^9,11,13,14^. However, the pharmacokinetics, safety, and efficacy of milrinone may vary with developmental stage, particularly in the immature pediatric population^11,15–21^. Despite its widespread use in pediatric patients, much of the evidence guiding milrinone dosing and clinical use is derived from studies in adults or heterogeneous pediatric populations.

Our recent work demonstrated age-dependent adaptations in cardiac electrophysiology and cardiac gene expression profiles, with the most pronounced changes occurring within the first year of life ^22–24^. However, limited access to pediatric cardiac tissue remains a major barrier to biomedical research, leaving much of our understanding of postnatal cardiomyocyte maturation and developmental adaptations reliant on animal models^25,26^. Animal models have also provided important insights into age-dependent responses to cardiac pharmacotherapies^27–30^. Thus, preclinical models provide an important opportunity to investigate how developmental maturation of the myocardium may influence responses to cardiac medications, including milrinone. Consistent with guidance from the Cardiac Safety and Research Consortium, well-characterized animal models can help address knowledge gaps in pediatric cardiac research and inform the development and clinical evaluation of therapies for children^31^.

To address this gap, we investigated how postnatal maturation influences myocardial responsiveness to acute milrinone exposure using a guinea pig model. The guinea pig heart shares several key features with the human heart, including ionic currents, action potential morphology, calcium handling, and force–frequency relationships^32^, and offers strong translational relevance due to comparable pharmacologic responsiveness^33,34^ and more human-like PDE activity and isoform expression^35^. In this study, we compared neonatal, juvenile, and adult guinea pig hearts by assessing cardiac structure, calcium-handling gene expression, electrophysiology, ventricular pressure development, contractile reserve, and optical mapping of cardiac action potentials and calcium transients. We hypothesized that postnatal maturation of cardiomyocyte structure and calcium handling would contribute to developmental differences in myocardial responsiveness to milrinone. This study underscores the importance of accounting for developmental stage when evaluating cardiac physiology, drug efficacy, and safety, as age-related adaptations may influence myocardial responses and contribute to variability in clinical outcomes.

## METHODS

### Animal model

The Institutional Animal Care and Use Committee of the Children’s Research Institute approved all animal procedures, which align with the guidelines in the National Institutes of Health’s Guide for the Care and Use of Laboratory Animals and the United States Department of Agriculture Animal Welfare Act. Experiments were performed using male and female Dunkin-Hartley guinea pigs (Elm Hill Labs: Chelmsford, MA US; Charles River Laboratories: Quebec, CA US; HillTop Lab Animals Inc.: Scottsdale, PA US). Animals were housed in conventional acrylic cages within the research animal facility, following standard environmental conditions including a 12 h light/dark cycle, a temperature range of 18–25°C, and humidity levels maintained between 30 and 70%. To evaluate age-specific differences, guinea pigs (n=86) were categorized into three age groups: neonates (99±14.37 g, 0–2 days old), juveniles (131.3± 27.4 g, 4-10 days old), and adults (840.2± 124.2 g, >6 months old). Guinea pigs were anesthetized with 2–3% isoflurane until achieving a surgical depth, as determined by an ear pinch test. Following a thoracotomy, the heart was excised and animals were euthanized by exsanguination.

### Microarray Experiments

Cardiac tissue samples were submerged in RNAlater stabilization solution (Invitrogen, Waltham, MA), temporarily stored at 4°C, and then transferred to –80°C until RNA extraction. Total RNA was isolated from right atrial and left ventricular tissue (n=30, n=5 per age group per chamber, 5–30 mg) and processed using an RNeasy fibrous tissue kit with on-column DNase treatment (Qiagen, Germantown, MD). RNA concentration was determined using a NanoDrop spectrophotometer (ThermoFisher, Waltham, MA), and RNA quality was assessed using a Bioanalyzer 2100 (Agilent Technologies, Santa Clara, CA). Total RNA (250 ng) input was primed for the entire length of RNA, including both poly(A) and non-poly(A) mRNA and reverse transcribed to generate sense-stranded targets that were biotin-labeled using a GeneChip WT Plus Reagent kit, and then hybridized to GeneChip™ Guinea Pig Gene 1.0 ST Array (Applied Biosystems, Waltham, MA, Catalog #902244) for 16 h at 45°C. Microarrays were washed and stained on the Fluidics Station F450 and then scanned using an Affymetrix GeneChip Scanner (GCS 3000 7 G; ThermoFisher). Quality control data and gene expression data were evaluated using Transcriptome Analysis Console (Applied Biosystems).

### Histology

Left ventricular tissue was fixed with 10% neutral buffered formalin and paraffin-embedded. For quantitative analysis, tissue sections (4-5 μm) were stained with hematoxylin and eosin, imaged using the same microscopy settings, and imported into ImageJ software (http://rsb.info.nih.gov/ij). The length of the intercalated discs were measured, and cardiomyocyte size was measured using the intercalated discs to demarcate cell-cell junctions. Measurements were collected from n=5 randomly selected cells across n=2-5 different regions per slide; the mean value was reported for each tissue preparation.

### Excised Heart Experiments

The whole, intact heart was cannulated at the aorta and transferred to a temperature controlled (37°C), constant pressure (68-70 mmHg) Langendorff-perfusion system, as previously described ^26,36^. A modified Krebs-Henseleit buffer was used for the perfusate, containing (in mM) 118.0 NaCl, 3.3 KCl, 2.0 CaCl_2_, 1.2 MgSO_4_, 1.2 KH_2_PO_4_, 24.0 NaHCO_3_, 10.0 glucose, 2.0 sodium pyruvate, and 10.0 HEPES buffer. The perfusate was continuously bubbled with carbogen (95% O_2_, 5% CO_2_). For each study, the heart was equilibrated for 20 min before collecting baseline measurements. Thereafter, the perfusate media was supplemented with sequential concentrations of milrinone (10 or 100 μM) for 15 min before collecting measurements. Concentration selection was guided by previously published studies ^37,38^. Pseudo-electrocardiograms (ECGs) were acquired throughout the study using a PowerLab acquisition system and analyzed in LabChart (ADInstruments, Colorado Springs, CO).

In a subset of studies, cardiac mechanical function was assessed by inserting a deionized water-filled latex balloon into the left ventricle^39^. The volume of the balloon was adjusted to maintain a diastolic pressure of 5-10 mmHg. Pressure changes were measured using a pressure transducer (Harvard Apparatus, Holliston, MA) connected to a bridge amplifier (Warner Instruments, Hamden, CT). To control the beating rate, both dynamic pacing (S1) and extra-stimulus pacing (S1-S2) protocols were driven by a stimulator (MappingLabs LtD: Oxford UK) connected to a bipolar stimulation electrode positioned on the left ventricle (1-3 mA, 1-2 ms pulse width).

Signals were acquired using a PowerLab acquisition system and analyzed in LabChart. Left ventricular developed pressure (LVDP) was calculated as the difference between minimum diastolic and maximum systolic pressure. Contractility was calculated as the rate of pressure development during systole (dP/dt_max_), and lusitropy as the rate of relaxation (dP/dt_min_). Post-rest potentiation was calculated as the LVDP difference between the S1 and a delayed S2 beat (200, 250, 300 ms or first sinus beat) to measure how the myocardium responds to an extended pause.

In a subset of studies, heart preparations were mechanically arrested with 12 μM (-/-) blebbistatin^40,41^. Hearts were loaded with a calcium indicator dye (50 μg Rhod-2, AM; AAT Bioquest: California US), which was recirculated for 10 minutes. Subsequently, a potentiometric dye was loaded (62 μg RH237 AAT Bioquest). The epicardial surface was illuminated using high-powered LED light sources (Solis-525C, ThorLabs: Virginia US), equipped with excitation filters (535 ± 25 nm). Fluorescence signals from Rhod-2 were acquired using a bandpass filter (585 ± 20 nm), while those from RH237 were filtered with a long-pass filter (>710 nm). Image stacks were collected from the anterior epicardium using an Optical Mapping System (MappingLabs LtD, Oxford UK) equipped with two Prime BSI cameras (Teledyne Photometrics: Arizona US) at 400-800 frames per second. Optical signals were analyzed using custom software ^42,43^. Action potential duration (APD) was measured as the difference between activation time and specific repolarization phases (30, 50, and 70%). Calcium transient duration (CaD) was calculated as the difference between the maximum slope of the calcium transient upstroke (dF/dt_max_) to specific phases of decay (30, 50, and 70%).

### Exclusion criteria

Animals were excluded from analyses if ECG recordings showed abrupt heart rate changes or poor signal-to-noise ratio which prevented accurate measurements. Optical recordings were excluded if there was inadequate signal-to-noise ratio or motion artifact. Individual paced beats were excluded if capture was not achieved at the intended pacing rate. Since the intrinsic heart rate varied substantially with developmental stage, some hearts could not be reliably captured at all pacing rates, and these data were excluded from analyses at the corresponding pacing interval.

### Statistical Analysis

Descriptive results are reported as mean ± standard deviation. Group comparisons with repeated measurements (i.e., baseline, 10, 100 μM milrinone) were evaluated using mixed-effects analysis with Geisser-Greenhouse correction and Holm-Sidak comparisons testing. Independent group comparisons (i.e., neonate, juvenile, adult) were evaluated using Welch’s ANOVA with Dunnett’s comparisons testing. Statistical significance was defined as a p-value less than 0.05, which is denoted in each figure by an asterisk. For microarray analysis, differentially expressed genes were identified by ANOVA with a 0.1 false discovery rate and heat maps were generated using log_10_ transformation and Z-score normalization for each gene signal intensity.

## RESULTS

### Developmental adaptations in cardiomyocyte anatomy and calcium-handling genes

First, we evaluated structural and molecular maturation of the guinea pig heart across each age group. During postnatal heart development, cardiomyocytes transition from hyperplastic to hypertrophic growth – which coincides with structural maturation of cardiomyocytes and an overall increase in muscle growth^44–46^. Structural development of T-tubules and their alignment with the sarcoplasmic reticulum help to support efficient calcium-induced calcium release in adult cardiomyocytes, as well as their responsiveness to inotropic agents^47–50^. Guinea pig heart weight increased relative to body weight across development, with neonates displaying the largest heart-to-body weight ratio (**Figure 1A**). As illustrated in **Figure 1B,C**, younger cardiomyocytes were smaller with a reduced length-to-width ratio (neonates: 2.58, juvenile: 4.98) and greater nuclei-to-cell area, while adult cardiomyocytes were more elongated (length-to-width ratio: 8.02) with prominent intercalated disc structures and a smaller nuclei-to-cell area ratio (**Supplemental Figure 1**). Cardiac development was also associated with increased expression of calcium-handling genes, including *Cacna1c*, *Ryr*, *Camk2d*, and *S100a1* in adult myocardium (**Figure 1D**).

**Figure 1.**
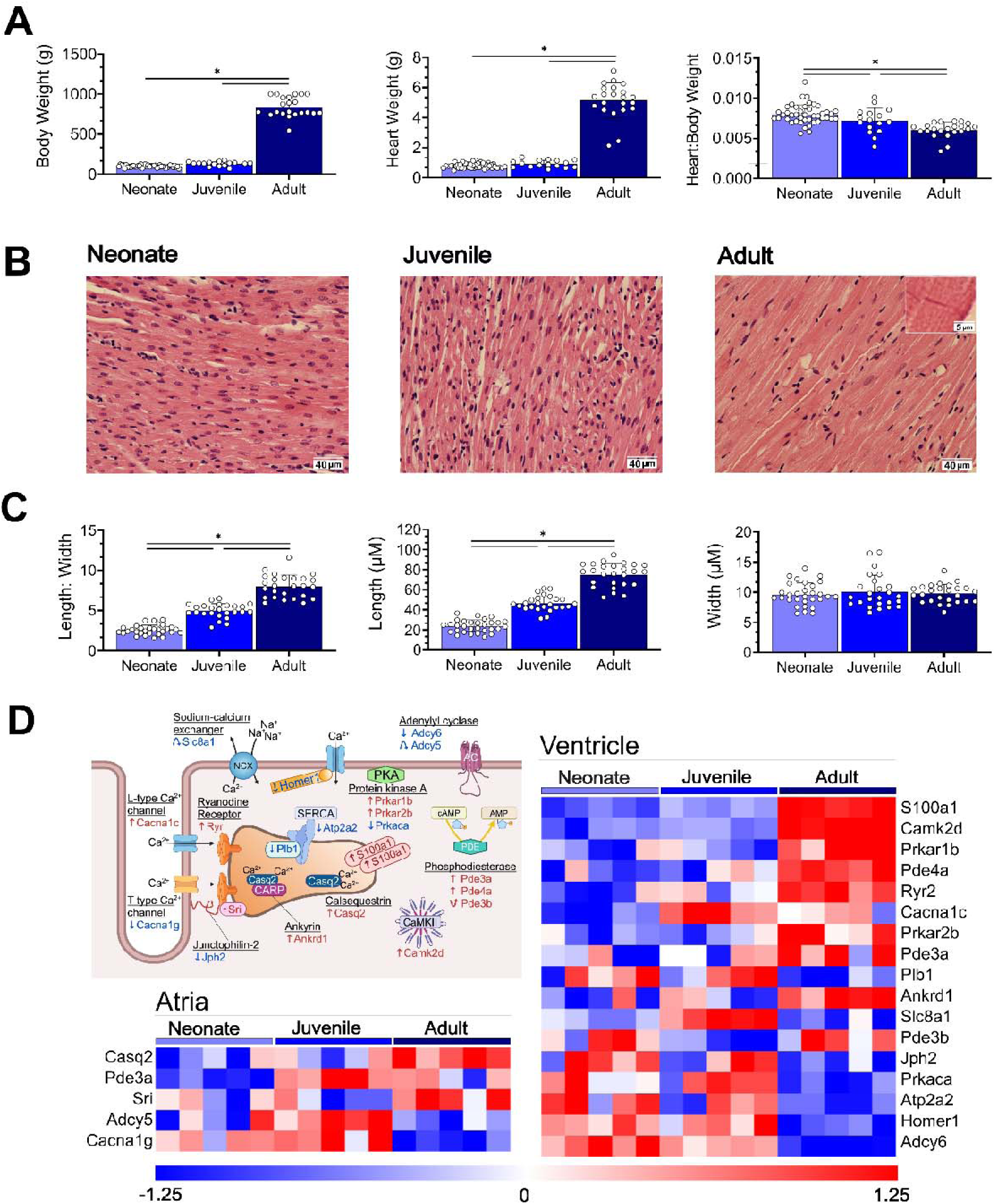
Developmental changes in cardiomyocyte structure and calcium handling genes. **A)** Relationship between heart and body weight during postnatal development. Neonates: 0-2 days (n=43), juveniles 4-10 days (n=16), adults (n=24). **B)** Examples of left ventricular heart tissue sections were stained with hematoxylin and eosin. **C)** Morphometric changes in cardiomyocyte width and length during postnatal development. Inset shows an intercalated disc in adult tissue section. Sample size includes multiple tissue sections for neonatal (N=6 hearts, n=32 sections), juvenile (N=5, n=25), and adult hearts (N=6, n=26). Data reported as the mean ± standard deviation, Welch’s ANOVA with Dunnett’s comparison testing, *p<0.05. **D)** Cartoon highlighting calcium handling genes. Heatmaps show differentially expressed genes involved in calcium handling (each row) for each animal (column; n=5 hearts per age group). On a per gene basis, the signal intensity was log_10_ transformed and Z-score normalized (blue: decreased, red: increased expression). Differentially expressed genes were identified by 1-way ANOVA with a 0.1 false discovery rate.

### Developmental differences in cardiac electrophysiology and the chronotropic response to milrinone

Having established developmental differences in cardiac structure and calcium-handling pathways, we next evaluated whether these adaptations were associated with differences in cardiac electrophysiology. Although milrinone is characterized as a PDE3 inhibitor with positive inotropic and vasodilatory effects, its effects on heart rate have varied across experimental models^51,52^. Prior studies have reported a half-maximal effective milrinone concentration ranging from 30-60 μM^37^. Accordingly, we tested the age-dependent effects of milrinone at a low (10 μM) and high (100 μM) concentrations on neonatal (0–2 days), juvenile (4–10 days), and adult isolated hearts (>6 months), with the two concentrations administered sequentially in the same heart. Under basal conditions, ex vivo ECG parameters differed across developmental stages (**Figure 2**), which is consistent with prior work^26,36^. Following milrinone administration, heart rate increased in all age groups and at both concentrations, with a greater intrinsic chronotropic response after escalation to 100 μM. At 10 and 100 μM, respectively, heart rate increased 24.0-33.5% (+62.7 to +87.4 bpm) in neonates, 18-35.3% (+42.5 to +83.4 bpm) in juveniles, and 19.5-35.1% (+37.9 to +68.3 bpm) in adults. Heart rate acceleration was accompanied by shortening of the QT interval, particularly at the 100 μM concentration (−39.6 ms neonates, −38.5 ms juveniles, −44.9 ms adults). P duration, PR interval, and the QRS interval were not significantly altered by milrinone.

**Figure 2.**
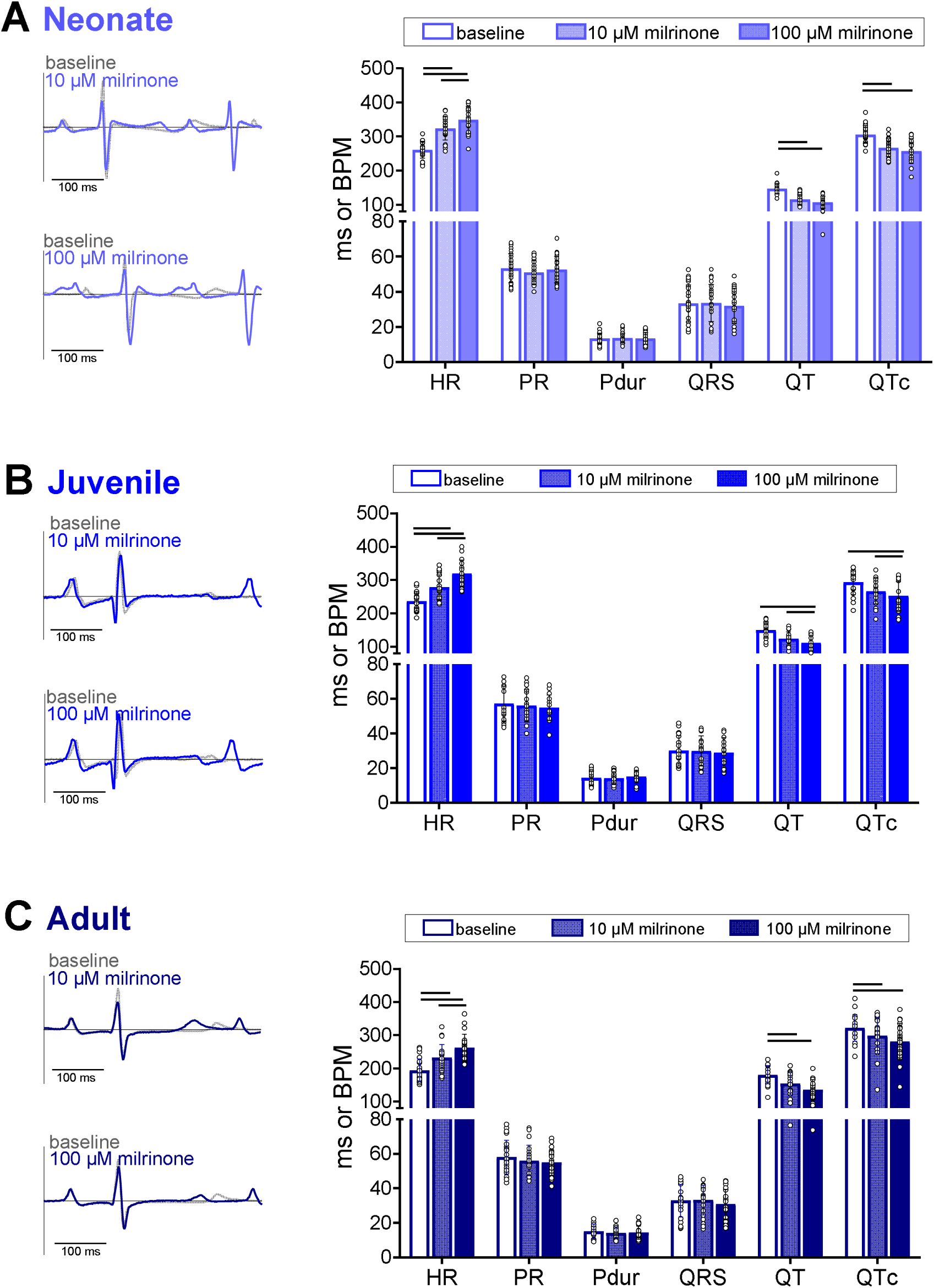
Developmental differences in cardiac electrophysiology and the chronotropic response to milrinone. Pseudo-ECGs were recorded from excised, intact heart preparations from **A)** neonates (n=22-26), **B)** juveniles (n=17-18), **C)** adults (n=21-22). Left: Representative ECG trace from guinea pig heart before and after milrinone treatment (ex vivo, retrograde perfusion). Right: ECG metrics for each age group before and after milrinone treatment. Individual replicates are shown; values are reported as mean ± SD. Comparisons before and after drug exposure by mixed effects analysis with Holm-Sidak multiple comparisons test; *p<0.05.

### Milrinone enhances contractility and relaxation across all developmental stages, with attenuated effects during rapid pacing

We next evaluated the effects of developmental stage and milrinone exposure on ventricular mechanical function. Guinea pig hearts exhibited a biphasic frequency–force relationship, with LVDP increasing during sinus acceleration (positive force–frequency relationship) but declining during external pacing at progressively shorter cycle lengths (negative force–frequency relationship; **Figure 3A,B**). Milrinone increased LVDP in hearts from all developmental stages, with the largest effects observed during sinus rhythm. Adult hearts exhibited the largest increase in LVDP (10 μM: +29 mmHg) and greatest increase in the positive LVDP slope during sinus acceleration (10 μM: 1.03 mmHg/BPM), whereas neonatal and juvenile hearts showed smaller but consistent responses (LVDP: +17.3 and +18.7 mmHg; slope: 0.40 and 0.65 mmHg/BPM, respectively at 10 μM). The increase in LVDP persisted during external pacing, but was attenuated at faster rates (**Figure 3C**). Although 10 and 100 μM concentrations were added sequentially, the mechanical response to milrinone did not follow the same concentration response relationship we observed with heart rate. In many hearts, the largest increase in LVDP occurred transiently following 10 μM milrinone, with LVDP partially returning toward baseline, and the subsequent response to 100 μM was often similar in magnitude.

**Figure 3.**
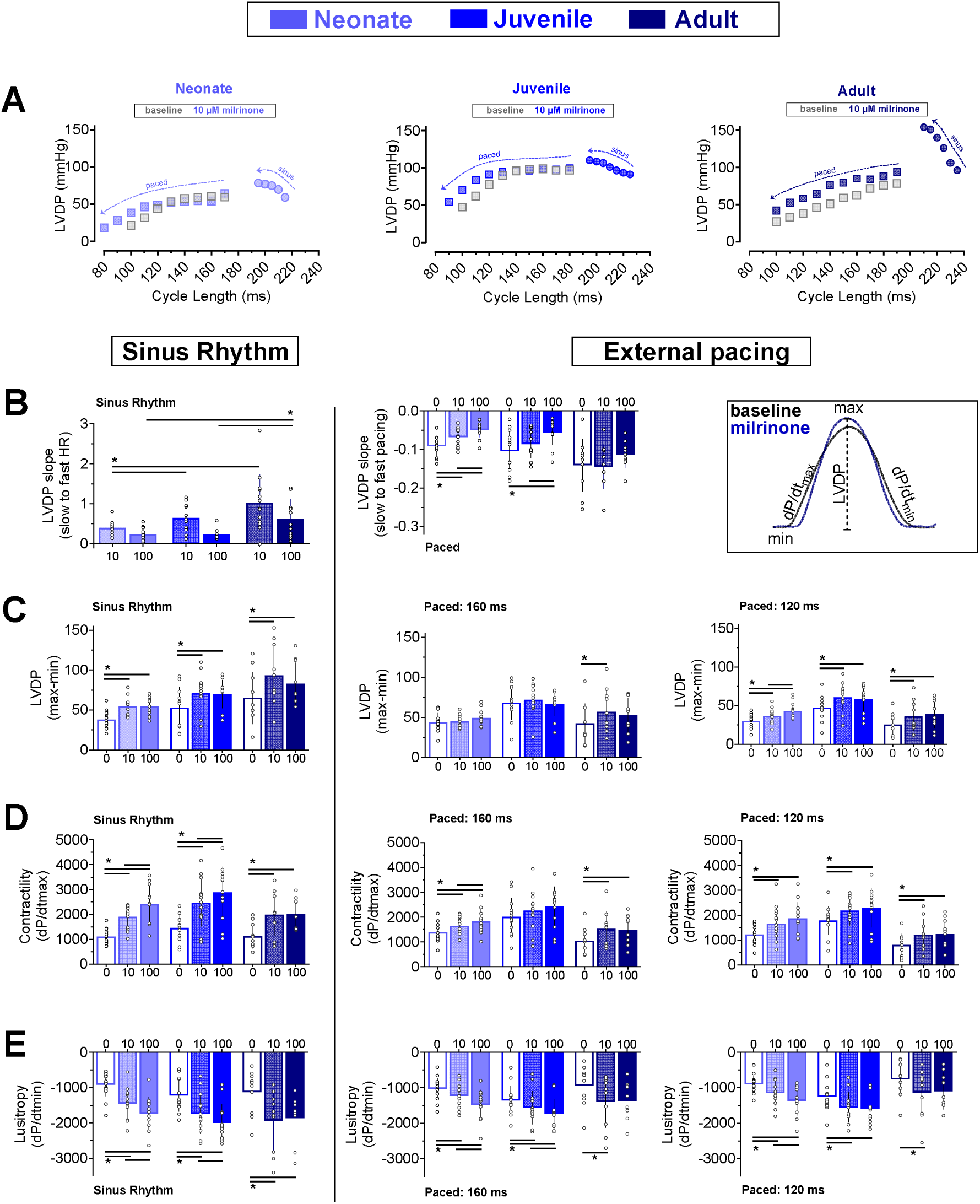
Milrinone enhances contractility and relaxation. **A)** Example of biphasic frequency response shown for a neonate, juvenile, and adult heart. LVDP increases with sinus rate acceleration (shorter cycle length), and decreases at increasingly faster pacing rates. **B)** Positive LVDP vs heart rate slope changes during accelerated sinus rhythm (left), vs negative slope with external pacing (center). Illustration of LVDP, contractility, and lusitropy measurements (right). Effects of milrinone on LVDP **(C)**, contractility **(D)**, lusitropy **(E)** are shown during sinus rhythm (left) and external pacing (right). Data represented as mean ± SD. Each dot represents an individual animal (n=14-15 neonates, n=13 juveniles, n=10-13 adults). Mixed effects analysis with Holm-Sidak multiple comparisons testing; *p<0.05.

Milrinone also enhanced indices of contraction and relaxation, with notable developmental differences in the magnitude and concentration-dependent responses (**Figure 3D,E**). During sinus rhythm, increases in contractility (dP/dt_max) were evident across all ages, with neonatal and juvenile hearts demonstrating a clear concentration-dependent response, whereas adult hearts exhibited a more modest response at the higher concentration (neonatal: +743 and +1,313 mmHg/s; juvenile: +1,010 and +1,425 mmHg/s; adult: +852 and +893 mmHg/s at 10 and 100 μM, respectively). Enhanced relaxation (dP/dt_min) was also evident across all age groups, although neonatal and juvenile hearts demonstrated a milrinone concentration-dependent response, while adult hearts exhibited a maximal response at 10 μM with no additional increase at 100 μM (neonatal: Δ543 and Δ812 mmHg/s; juvenile: Δ513 and Δ775 mmHg/s; adult: Δ806 and Δ733 mmHg/s at 10 and 100 μM, respectively). These effects persisted during external pacing, but were attenuated at faster rates (**Figure 3D,E**).

### Post-rest potentiation increases with maturation and is reduced after milrinone exposure

We next examined post-rest potentiation as an additional measure of contractile reserve. Post-rest potentiation is a phenomenon in which a brief rest interval enhances the force of the subsequent contraction through increased calcium loading and release. Post-rest potentiation was greater in adult than neonatal hearts across the S1–S2 intervals tested, with juvenile hearts showing an intermediate response (**Figure 4**). At baseline, adult ΔLVDP increased progressively with longer rest intervals (200 ms: 34.9±16, 250 ms: 59.5±17.3, 300 ms: 76.1±17.7 mmHg), whereas neonatal responses were smaller and less dependent on the rest duration (200 ms: 11.8±5.8, 250 ms: 18.1±8.7, 300 ms: 20.4±9.8 mmHg). This difference was also evident when comparing the first spontaneous contraction with the preceding beat (adult: 94.6±38.5, neonate: 28.3±13.1 mmHg; **Figure 4G,H**). Milrinone reduced post-rest potentiation across all developmental stages. In neonatal hearts, the increase in contractile performance left relatively little additional force to be recruited after rest. Adult hearts retained post-rest potentiation after 10 μM milrinone, but the response was further reduced after escalation to 100 μM. Thus, the reduction in post-rest potentiation after milrinone is therefore consistent with increased baseline performance and a reduced capacity for further force augmentation following rest, rather than necessarily indicating impaired calcium storage or release.

**Figure 4.**
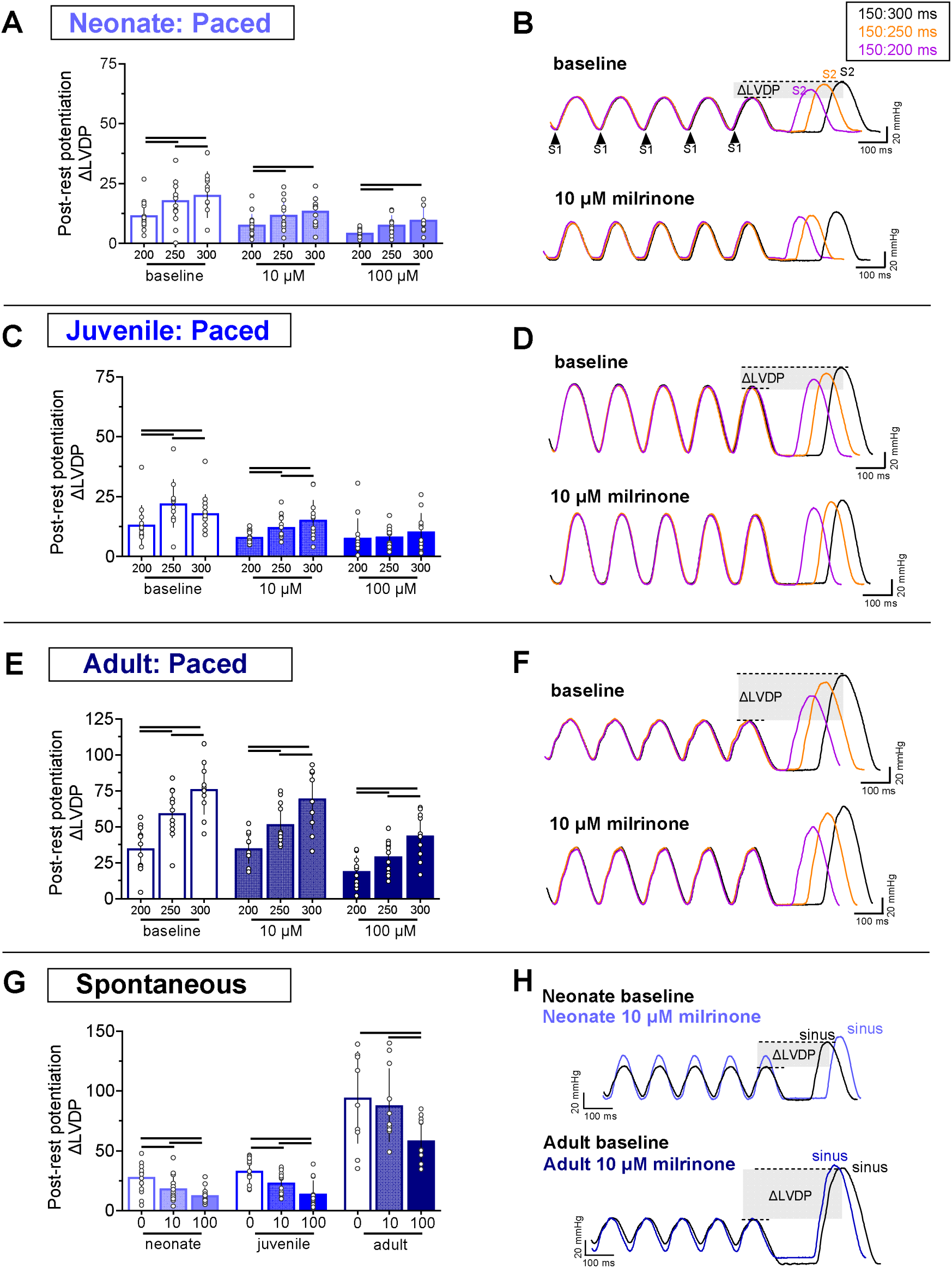
Post-rest potentiation increases with maturation and is reduced after milrinone exposure. S1-S2 pacing protocols were implemented to measure post-rest potentiation with across multiple rest periods (300 ms, 250 ms, 200 ms), and the corresponding change in LVDP is shown for **A,B)** neonates, **C,D)** Juveniles, **E,F)** adult hearts. For each age group the compiled data is shown (left), with representative pressure wave examples (right). Dynamic pacing is shown (S1 carrots, 150 ms), followed by the S2 beat, and the change in LVDP is shown. **G,H)** Change in LVDP from S1 dynamic pacing (120 ms) to the first spontaneous beat. Data represented as mean ± SD. Each dot represents an individual animal (n=11-16 neonates, n=12-13 juveniles, n=10-11 adults). Mixed effects analysis with Holm-Sidak multiple comparisons testing; *p<0.05.

### Milrinone differentially modulates cardiac repolarization and calcium transient kinetics across development

Although milrinone consistently accelerated heart rate and enhanced mechanical performance, its effects on cardiac electrical activity and calcium dynamics remained unclear. We therefore used optical mapping to measure action potential duration (APD) and calcium transient duration (CaD) during dynamic pacing, allowing these responses to be evaluated independently of changes in sinus rate (**Figure 5,6**). Across all developmental ages, APD displayed rate-dependent restitution kinetics for each repolarization phase measured (APD_30_, APD_50_, APD_70_; **Figure 5A**). APD was longest in adults, intermediate in juveniles, and shortest in neonates; at a 200 ms pacing cycle length, APD_50_ was 112.7±9.4 ms in adults, 93.1±14.5 ms in juveniles, and 87.5±9.6 ms in neonates. Milrinone did not significantly alter the slope of the APD restitution curves, indicating that the fundamental rate dependence of repolarization was preserved (**Figure 5B,C**). Milrinone shortened APD measurements in neonates and adults; at 200 ms pacing cycle length, APD_50_ shortened by 15.3-27.1% in neonates (10 μM: 74.1±13.7, 100 μM: 63.8±12.9 ms) and 8.6-15.2% in adults (10 μM: 103±3.3, 100 μM: 95.6±3.9 ms), but the effects on juvenile hearts were minimal and not significant (**Figure 5D**).

**Figure 5.**
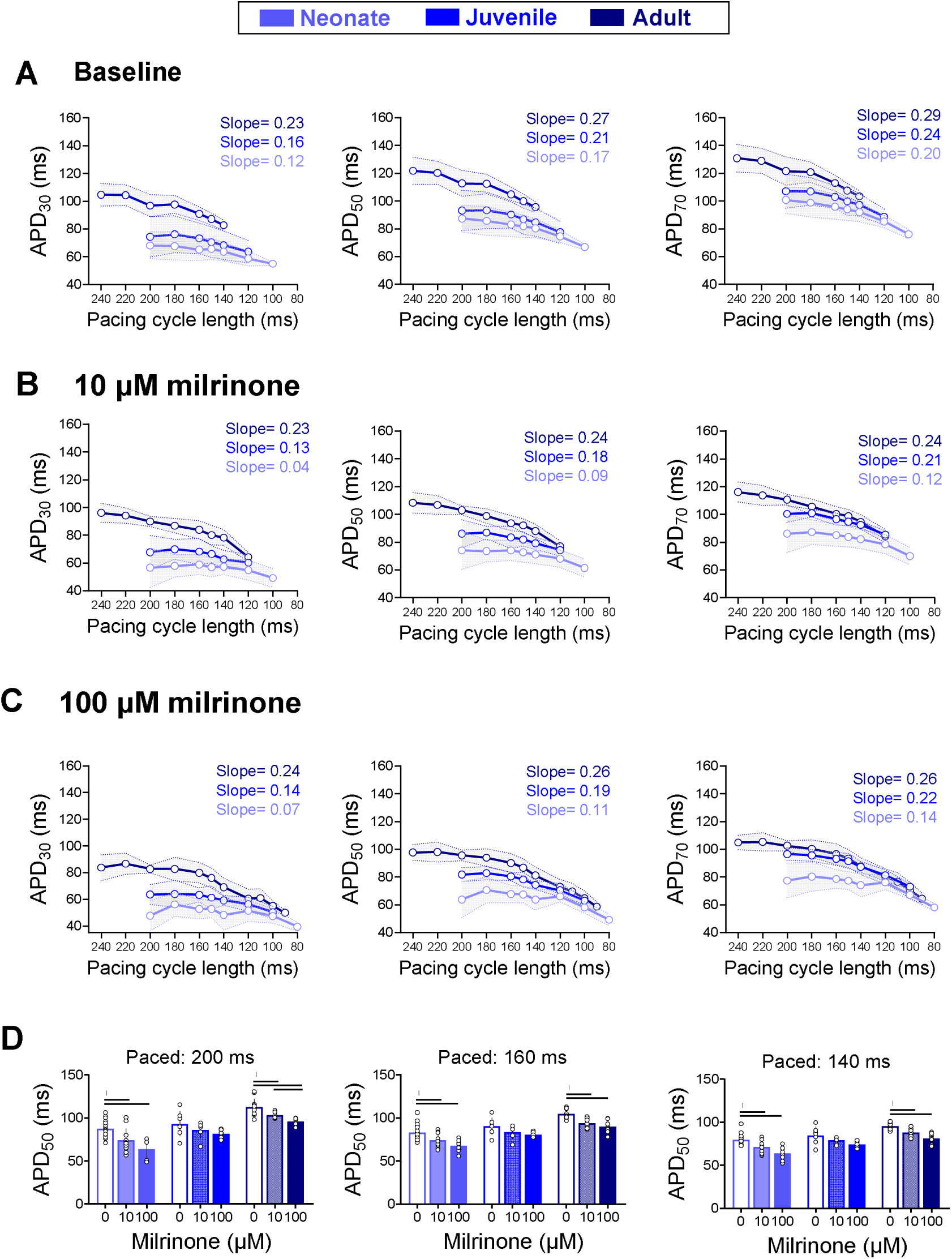
Effects of milrinone on cardiac action potential duration. Optical action potential duration measurements are plotted relative to pacing cycle lengths at **A)** baseline, **B)** after 10 μM milrinone treatment, **C)** after 100 μM milrinone treatment. Slope of the restitution curve is shown for each age group **D)** Comparison between baseline and milrinone measurements at APD50 are shown for 200 ms, 160 ms, and 140 ms pacing cycle length. APD30 = action potential duration at 30% repolarization, APD50 = 50% repolarization, APD70 = 70% repolarization. Data represented as mean ± SD (shown as a line in restitution curves and error bar in bar plot). Each dot represents an individual animal (n=5-19 neonates, n=6-7 juveniles, n=8-13 adults). Mixed effects analysis with Holm-Sidak multiple comparisons testing; *p<0.05.

**Figure 6.**
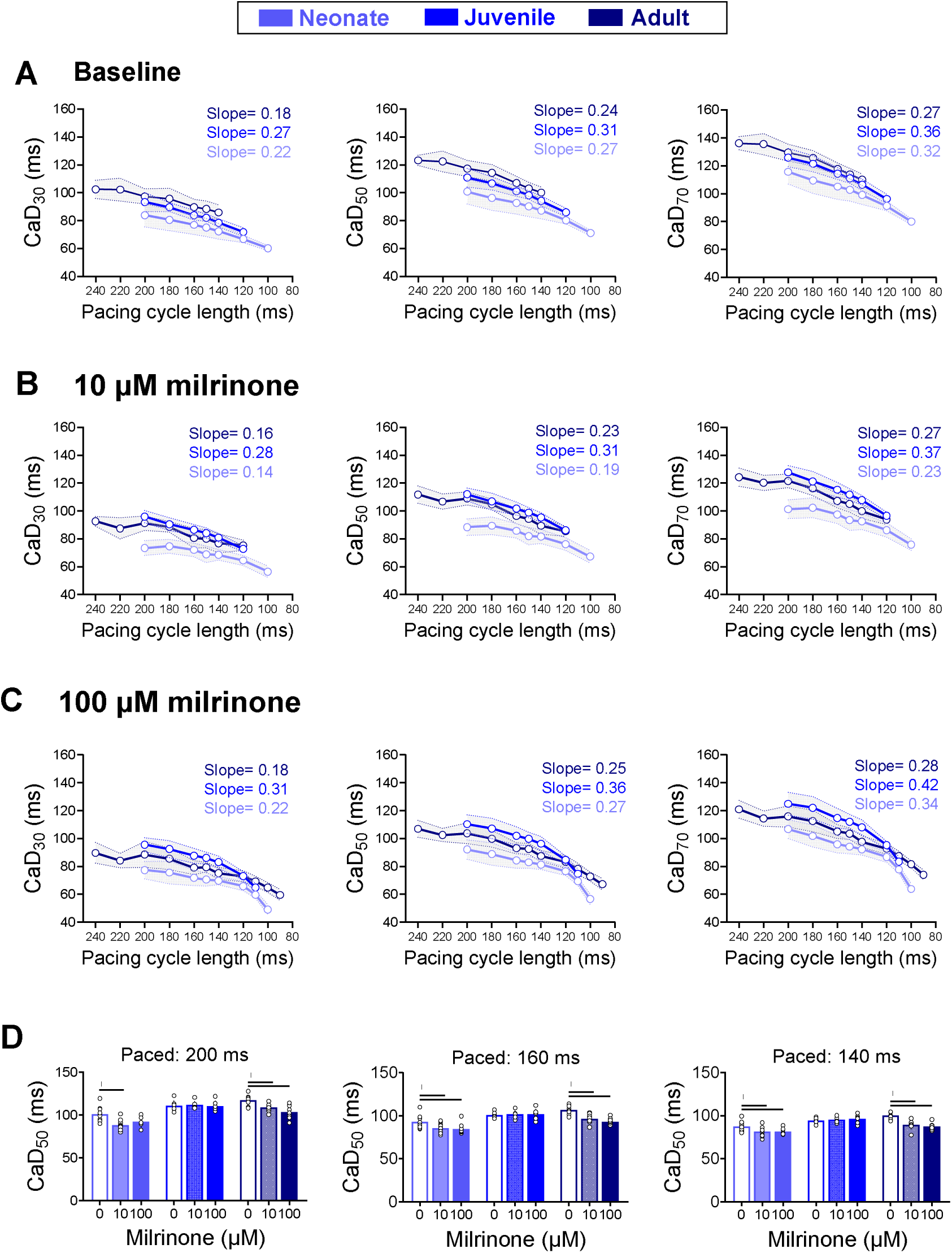
Effects of milrinone on intracellular calcium handling. Optical calcium transient measurements are plotted relative to pacing cycle lengths at **A)** baseline, **B)** after 10 μM milrinone treatment, **C)** after 100 μM milrinone treatment. Slope of the restitution curve is shown for each age group. **D)** Comparison between baseline and milrinone measurements at CaD50 are shown for 200 ms, 160 ms, and 140 ms pacing cycle length. CaD30 = calcium transient duration at 30% decay, CaD50 = 50% decay, CaD70 = 70% decay. Data represented as mean ± SD (shown as a line in restitution curves and error bar in bar plot). Each dot represents an individual animal (n=5-11 neonates, n=6 juveniles, n=8-10 adults). Mixed effects analysis with Holm-Sidak multiple comparisons testing; *p<0.05.

Given the developmental differences we observed in mechanical function and post-rest potentiation, we next evaluated age-dependent changes in intracellular calcium handling. Calcium transient duration (CaD_30_, CaD_50_, CaD_70_) also displayed rate dependent kinetics, with shorter calcium transients at faster pacing rates (**Figure 6A**). Baseline CaD values were longer in adults and juvenile hearts (CaD_50_: 117.4±5.8 ms adult, 110.9±6.7ms juvenile) and shorter in neonatal hearts (CaD_50_: 100.8±8.8 ms at 200 ms pacing cycle length). Milrinone shortened CaD_50_ by 8.6-12.7% in neonatal hearts (10 μM: 88.4±6, 100 μM: 92.1±7 ms) and 7.2-11.7% in adult hearts (10 μM: 108.9±4.8, 100 μM: 103.7±7.5 ms, 200 ms pacing), but had no measurable effect in juveniles (**Figure 6B-D**). Together, these findings demonstrate that milrinone produces developmentally distinct effects on cardiac electrical activity and calcium kinetics, with neonatal and adult hearts exhibiting shortening of both APD and CaD, whereas juvenile hearts were largely unresponsive.

## DISCUSSION

In this study, we examined how postnatal maturation influences the acute myocardial response to milrinone using an intact guinea pig heart model. Collectively, our main findings include: (i) neonatal guinea pig hearts display distinct expression of calcium-handling genes compared to juveniles and adults, (ii) milrinone produced chronotropic effects across all age groups, demonstrating that PDE3 inhibition directly accelerates the intrinsic heart rate, (iii) milrinone enhanced ventricular contractility and relaxation at every age, but the magnitude and frequency-dependence of this response varied with development – with adults showing the greatest increase in LVDP during sinus rhythm and largest post-rest potentiation, (iv) despite robust mechanical effects, the effect of milrinone on APD or CaD shortening was modest and limited to neonates and adults, with little effect on juveniles.

### Developmental maturation of cardiac structure and signaling

Developmental changes in cardiac growth, cardiomyocyte structure, and calcium-handling gene expression provide an important substrate for physiological differences observed across age. Consistent with previously published allometric relationships between cardiovascular variables and body mass^53^, we observed a strong relationship between heart and body weight across neonatal, juvenile, and adult animals. This pattern parallels human cardiac growth, in which cardiac mass increases with body size during early development before relative cardiac weight stabilizes during later maturation^54,55^. Related to cardiac growth, the transition from cardiomyocyte hyperplasia to hypertrophy is a defining feature of postnatal cardiac maturation across species. In the present study, cardiomyocyte length increased 3.1-fold from the neonatal to adult period, consistent with prior observations in humans demonstrating a 3-fold increase in cardiomyocyte diameter between birth and adulthood^54,55^. This structural maturation occurs alongside progressive development of the excitation–contraction coupling machinery. We observed age-dependent increases in expression of genes involved in mature calcium-induced calcium release, including L-type calcium channels (*Cacna1c*) which localize to T-tubules near ryanodine receptors (*Ryr*) on the junctional sarcoplasmic reticulum (regulated by *S100a1*). These findings are consistent with both human and rodent studies reporting developmental increases in Cacna1c and Ryr expression and maturation of intracellular calcium handling^24,25,56^. The reduced post-rest potentiation observed in neonatal hearts is a reflection of a less mature sarcoplasmic reticulum-dependent reserve, consistent with the progressive maturation of calcium cycle observed at the molecular level.

Milrinone exerts its primary cardiac effects through inhibition of phosphodiesterase 3 (PDE3), thereby increasing cAMP signaling and enhancing myocardial contractility and relaxation. The developmental increase in *Pde3* expression observed in the present study, including *Pde3a*, raises the likelihood that maturation of the PDE3–cAMP signaling axis contributes to the greater mechanical response observed in adult heart preparations. Indeed, previous studies have demonstrated that Pde3a-deficient hearts exhibit enhanced basal contractility and relaxation, and are also unresponsive to additional PDE3 inhibition by milrinone^57^. The importance of age-dependent PDE regulation is further supported by studies using human myocardial specimens. Specifically, in the pediatric myocardium, chronic PDE3 inhibition has been associated with increased cAMP and phospholamban phosphorylation – while the adult myocardium demonstrated increased total and PDE3-specific activity, without a corresponding increase in cAMP or phospholamban phosphorylation^16^. Further, pediatric and adult failing hearts have been shown to have distinct age- and disease-associated expression levels of adenylyl cyclase and PDE isoforms, with differential responses to PDE3 inhibition^17^. Collectively, these findings suggest that developmental differences in cAMP regulation may influence myocardial responsiveness to milrinone, as observed in the present study. However, additional work is needed to fully elucidate the developmental trajectory of this signaling pathway, including direct assessment of PDE3A protein abundance, enzymatic activity, cAMP concentrations and compartmentalization, and downstream phosphorylation – which is beyond the scope of the current study.

### Developmental differences in the chronotropic and electrophysiological response to milrinone

One of the most consistent findings in the present study was the increase in sinus rate after milrinone exposure at every developmental stage. This finding is notable, as milrinone is generally introduced clinically as an inodilator, whose principal myocardial action is positive inotropy^9,11^ mediated by PDE3 inhibition, rather than direct stimulation of beta-adrenergic receptors. However, PDE3 inhibition increases intracellular cAMP by reducing its degradation, and cAMP signaling can influence pacemaker activity and calcium currents^58^. Prior experimental studies have documented positive chronotropic effects of milrinone in mammalian models^52^. The present study extends these observations across postnatal development and demonstrates that the chronotropic response is intrinsic to the isolated heart, rather than dependent on sympathetic activation. The chronotropic effects of milrinone have important clinical implications, given the recognized association between milrinone therapy and tachyarrhythmias. Indeed, clinical observations indicate that pediatric patients receiving milrinone following congenital heart surgery have increased odds of developing postoperative tachyarrhythmias^8,59^. Importantly, milrinone-associated tachyarrhythmias can have clinically significant consequences, including impaired myocardial perfusion and hemodynamic instability.

In addition to its chronotropic effects, milrinone altered ventricular repolarization in a developmental- and concentration-dependent manner. ECG recordings demonstrated concentration-dependent shortening of the QT interval, consistent with a faster heart rate because QT is strongly rate-dependent. However, significant shortening of Bazett-corrected QTc was also observed, suggesting that the effect may not be solely attributed to an elevated heart rate. This interpretation was partially supported by modest APD shortening in neonatal and adult hearts during external pacing, indicating that the effect on ventricular repolarization persisted when rate was controlled. Our observations of APD shortening are consistent with prior reports, including APD shortening in isolated canine ventricular muscle^60^ and isolated guinea pig ventricular cardiomyocytes^38^ with greater effects observed at increasing milrinone concentrations. These results align with previous findings in which milrinone decreases the total duration of the action potential through more rapid repolarization^61^. The mechanisms underlying milrinone-induced APD shortening remain incompletely defined. PDE3 inhibition increases intracellular cAMP, which can modulate multiple ion channels involved in cardiac excitation and repolarization. Although cAMP-dependent enhancement of L-type calcium current (I_Ca,L) has been described^62,63^, this mechanism would extend the action potential plateau phase rather than accelerating repolarization. Thus, the net shortening of APD likely reflects the integrated effects of altered cAMP signaling on multiple depolarizing and repolarizing currents, rather than a single ionic mechanism. Developmental differences in these currents may contribute to the age-specific responses observed in the present study. In support of this possibility, L-type calcium current density has been reported to increase with maturation in guinea pig ventricular myocytes^64^, while differences in L-type calcium channel function have also been described between human infant and adult atrial cardiomyocytes^65^. Notably, in the present study, juvenile guinea pig hearts exhibited a comparable chronotropic response to neonates and adults – but, without significant APD shortening. This further suggests that developmental remodeling of cardiomyocyte anatomy, ionic currents, and cAMP signaling pathways influence myocardial responsiveness to milrinone. Collectively, these findings emphasize that the effects of milrinone extend beyond its well-characterized effects on myocardial contractility and vary across developmental stage.

### Effects of milrinone on contractile function, post-rest potentiation, and calcium handling

Prior human studies have described developmental differences in the cardiac force-frequency relationship, wherein shortening the pacing cycle length increased developed force in infant but not neonatal ventricular muscle strips^66^. Similarly, guinea pig papillary muscle demonstrated maximal force of contraction after milrinone treatment at a faster stimulation rate than under control conditions^61^. In agreement with previous studies^67,68^, we observed a biphasic force-frequency relationship across all developmental stages, with force increasing with an accelerated sinus rate to a maximum and then declining at faster (external) pacing frequencies. Although milrinone maintained a positive mechanical effect across multiple pacing frequencies, the magnitude of this response was attenuated at faster rates, suggesting that the effects of PDE3 inhibition interact with the intrinsic frequency dependence of calcium cycling and diastolic relaxation. In our study, we also observed age-dependent differences in post-rest potentiation, with adult hearts demonstrating the greatest left ventricular pressure following a period of rest. Post-rest potentiation reflects progressive calcium loading into the sarcoplasmic reticulum (SR) during rest, and thus, is more pronounced in the mature myocardium with a well-developed SR^69,70^. Thus, the reduced post-rest potentiation observed in neonatal hearts is consistent with the immature SR phenotype and developmental differences in calcium-handling gene expression observed in the present study. The reduction in post-rest potentiation following milrinone, however, should not necessarily be interpreted as impaired contractile reserve, as the increased baseline performance produced by milrinone leaves less additional force to recruit following a rest interval. The intrinsically higher heart rate of neonatal hearts may also contribute to their smaller post-rest response.

Our calcium imaging data provide a complementary perspective on developmental differences in mechanical function. At baseline, neonatal hearts exhibited shorter calcium transient duration (CaD) than juvenile and adult hearts, consistent with immature SR calcium storage and a greater reliance on trans-sarcolemmal calcium influx during early postnatal development^71^. Following milrinone administration, neonatal and adult hearts demonstrated the greatest CaD shortening across pacing frequencies and concentrations, while juvenile hearts were largely unaffected. These findings are consistent with studies demonstrating that milrinone can accelerate calcium transient decay in human ventricular myocytes in a concentration-dependent manner ^72^, with evidence that increased cAMP signaling can enhance the rate of SR calcium uptake^73^. Accelerated calcium clearance can therefore contribute to the enhanced lusitropic response to milrinone. Importantly, our studies measured calcium transient duration rather than absolute intracellular calcium concentration or SR calcium content, and thus, CaD shortening should be interpreted as an alteration in calcium kinetics. Developmental differences in cAMP signaling, calcium cycling, or myofilament responsiveness are likely to contribute to the age-dependent mechanical response observed in the present study.

Taken together, our results suggest that developmental maturation changes the myocardial context in which PDE3 inhibition operates rather than producing a more simplistic linear age-dependent increase or decrease response to milrinone. Adult hearts had the greatest mechanical response and most mature post-rest reserve, while neonatal hearts exhibited less SR-dependent contractile reserve but maintained a pronounced heart rate response to milrinone. Juvenile heart responses were particularly interesting for their relative resistance to APD and CaD shortening, despite clear positive chronotropic and inotropic responses to milrinone. These results suggest that the multiple pathways governing mechanical performance and electrical / calcium duration are related, but not interchangeable across age groups. A developmental shift in PDE3 expression, calcium-handling machinery, cAMP compartmentalization, and myofilament sensitivity could collectively account for these varied phenotypes.

## CONCLUSIONS

Milrinone is among the most prescribed inotropic agents for increasing cardiac output during the perioperative and postoperative management of pediatric patients, yet dosing practices remain highly variable among providers and institutions^74^. This variability is particularly relevant given that the physiological response to milrinone may be influenced by developmental stage and underlying cardiac disease, while treatment has been associated with adverse effects including tachycardia, unpredictable concentration-response relationships, and tissue hypoxia^75–77^. However, limited clinical data are available to determine how developmental age or disease influences myocardial responses to milrinone

Using a translational guinea pig model of postnatal cardiac development, our study demonstrates that the physiological effects of milrinone are age- and concentration-dependent and are influenced by developmental changes in cardiac structure, calcium handling, and electrophysiology. These findings extend our previous work demonstrating progressive electromechanical remodeling of the guinea pig heart during postnatal development^26^ and provide evidence that developmental maturation influences the myocardial response to PDE3 inhibition. Collectively, these findings highlight the importance of considering developmental stage when evaluating milrinone pharmacology in pediatric patients and underscore the need for further investigation of age- and concentration-dependent responses to milrinone and their relationship to clinical outcomes following pediatric cardiac surgery.

## LIMITATIONS

Experiments were performed in isolated Langendorff-perfused hearts, which removes autonomic and hemodynamic influences. This limitation is particularly relevant to the interpretation of the chronotropic response and limits direct extrapolation to postoperative patients. Further, our study utilized two milrinone concentrations that were administered sequentially to the same heart; therefore, the 100 μM condition represents escalation rather than an independent group. The experimental concentrations used in the isolated heart preparations should not be equated directly with clinical plasma concentrations or infusion rates. We also report gene expression measurements – yet additional studies are necessary to identify any corresponding changes in protein abundance, enzymatic activity, cAMP compartmentalization, or phosphorylation of downstream calcium handling proteins. Further, we used fluorescence-based measurements of voltage and calcium kinetics rather than measurements of absolute membrane potential or absolute intracellular calcium concentrations. Finally, guinea pig hearts provide important translational advantages but retain species-specific differences from the human myocardium and LV balloon-derived pressure measurements are a surrogate for direct measurements of in vivo cardiac output.

## Acknowledgements

We gratefully acknowledge Dr. Susan Knoblach and the Children’s National Genomics Core for scientific and technical guidance related to guinea pig microarray gene expression experiments.

## Data Availability

Derived data supporting the findings of this study are available in this study, or specific datasets can be provided from the corresponding author (NGP) upon request.

## Supplement

**Supplemental Figure 1.**
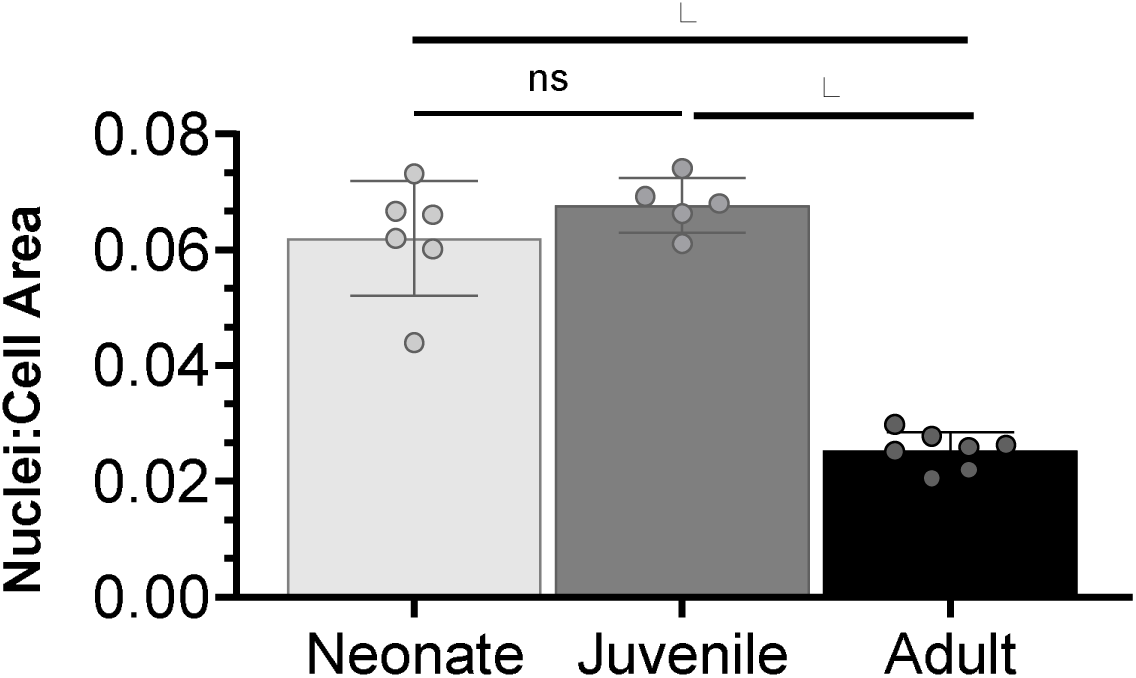
Cardiomyocyte size increases with development. Tissue sections from neonatal, juvenile, and adult hearts (n=5-7) were stained with hematoxylin and eosin, imaged, and the total nuclei area was measured relative to the cell area in each field of view. Adult cardiomyocytes had a smaller nuclei-to-cell area, indicative of larger cardiomyocytes in hypertrophic growth phase. Data reported as the mean ± standard deviation*p<0.05.

